# Transcriptomic assessment of the effects of cisplatin and TiO₂ nanoparticles in Drosophila melanogaster

**DOI:** 10.1101/2025.11.03.685613

**Authors:** C. N. Costa, L. P. Ciapina, A. C. Bahia, V. Braz, F. A. B. Da Silva, F. Lopes

## Abstract

Cisplatin, the first metal-based chemotherapeutic drug, remains widely used despite its toxicity. Combining cisplatin with nanoparticles has been proposed to improve its therapeutic profile, although most studies rely on *in vitro* models. Using RNA-seq and bioinformatic analyses, we investigated the *in vivo* transcriptional effects of cisplatin (50 μg/ml) and titanium dioxide nanoparticles (TiO_2_ NPs; 50 μg/ml), alone and in combination, in *Drosophila melanogaster*. Flies exposed to cisplatin or TiO_2_ NPs alone exhibited modulation of genes associated with xenobiotic metabolism and detoxification. In contrast, combined exposure resulted in a markedly reduced number of upregulated differentially expressed genes (DEGs): CG10013, aqz, CG5568, and CG3213, which are related to cell division and DNA/RNA metabolism, likely due to a synergistic effect of the cisplatin and TiO2 administered simultaneously. We also observed that combined exposure promotes downregulation of genes involved in xenobiotic metabolism, detoxification, innate immune collapse, and reproduction and fertility. Although mortality rates were not significantly affected in any group, flies from the CIS/NPTIO2 group exhibited impaired climbing performance. These results suggest that co-exposure to cisplatin and TiO_2_ nanoparticles induces transcriptional suppression, potentially impairing essential cellular processes.

## INTRODUCTION

Cisplatin is the commercial name for the compound (SP-4-2)-diamminedichloridoplatinum (II), the first metal-based anticancer drug widely used to treat several cancers (Mohammadi; Li; Ewing, 2018; Rosemberg *et al*., 1969; Zhang *et al*., 2021). Multiple *in vitro* and *in vivo* studies show that its mechanism of action involves generating DNA lesions. These lesions trigger several signal transduction pathways that lead to inhibition of replication and transcription, cell cycle arrest, DNA repair, and apoptosis. DNA damage results from cisplatin’s interaction with purine bases in DNA, generating interstrand adducts (G-G, 60%; A-G, 20%) and DNA-protein interactions (Allgayer *et al*., 2019).

A genotoxic study using the Comet assay indicates that, in *Drosophila melanogaster*, detected DNA damage contains higher levels of G–G cross-links than G monoadducts, thereby contributing to DNA strand breaks by blocking DNA replication and repair (García Sar *et al*., 2012). Cisplatin is associated with side effects such as nephrotoxicity and ototoxicity in humans (Karasawa; Steyger, 2015), as well as chemotherapy-induced peripheral neuropathy (CIPN) in mammals and non-mammals, such as *D. melanogaster* (Groen *et al*., 2022).

Besides its side effects, cisplatin is still largely used as a chemotherapeutic drug. To reduce cisplatin’s side effects, research has examined the use of different carriers in combination with cisplatin. In this context, the combined use of 50 µg/ml TiO_2_ nanoparticles and 10 µg/ml cisplatin improved chemotherapy response *in vitro* in the F10 melanoma model and *in vivo* in C57BL/6 mice, likely through the crucial role of nanoparticle-induced autophagy in sensitizing F10 melanoma cells (Adibzadeh *et al*., 2021).

Although largely used in food, pharmaceutical, and cosmetic industries, titanium dioxide nanoparticles (NPTiO2) remain under investigation, with conflicting findings reported in the literature. While many authors describe *in vivo* toxic effects induced by TiO_2_ nanoparticles, such as altered lysosomal and oxidative stress biomarkers and reduced transcription levels of immune-related antioxidant genes (Barmo *et al*., 2013), as well as genotoxic effects like DNA damage and the induction of chronic gastritis (Mohamed, 2015). Other study that examined 34 robust datasets did not support a direct DNA-damaging mechanism for TiO_2_ in nano- or microform (Kirkland *et al*., 2022).

*Drosophila melanogaster*, commonly known as the fruit fly, has been used as a model organism for *in vivo* studies to elucidate many human diseases, like cancer (FREIRES *et al*., 2017), and substance toxicity (Groen *et al*., 2022).

Here, we investigate the *in vivo* effects of combined oral administration of cisplatin (50μg/ml) and TiO_2_ nanoparticles (50μg/ml), which may trigger distinct metabolic and molecular responses in *D. melanogaster*. To evaluate this, we analyzed gene modulation by identifying differentially expressed genes (DEGs) from RNA-seq data and performed downstream analysis to determine the ontology terms associated with the different treatments. Our *in vivo* results indicate that the combined use of cisplatin and TiO_2_ nanoparticles triggers a toxic effect, evidenced by a specific expression profile characterized by intermediate transcript levels and a minimal number of upregulated differentially expressed genes (DEGs).

## MATERIALS AND METHODS

### 1. *Drosophila melanogaster* maintenance

*Drosophila melanogaster* strain Oregon (RRID:BDSC.5) was obtained from the Bloomington Drosophila Stock Center, Indiana University. The culture was performed in a standard cornmeal-yeast-agar medium (BRENT; OSTER, 1974) and maintained in an incubator at 25 °C and humidity between 70% and 80%, with a 12-hour light cycle.

### 2. Treatments

1-to-4-day-old *D. melanogaster* strain Oregon flies were divided into four groups, each with 30 individuals. Each group contained a 4:1 female/male ratio. The control group and all treatment groups were performed in biological triplicate. The control group received a standard medium enriched with a mixture of yeast, water, and a green food coloring dye.

Treatment groups were treated with the same culture medium but with the following aqueous solutions instead of water: CIS group – aqueous cisplatin solution (50µg/ml); NPTIO2 group - aqueous NPTiO_2_ solution (50µg/ml); CIS/NPTIO2 group – cisplatin aqueous solution (50µg/ml) and NPTiO_2_ aqueous solution (50µg/ml) simultaneously. Flies were kept in this condition for 3 (three) days.

Only female flies were used for the RNA-Seq experiment.

### 3. Preparation of Used Solutions

NPTiO_2_ water solution was prepared using 50mg/ml of TiO_2_ (Sigma-Aldrich, Cat. No.: 718467; nano powder, 21 nm primary particle size (TEM), ≥99.5% trace metals basis), which was heated in an oven (200°C) for 30 minutes. Then, TiO_2_ was solubilized in deionized water by sonication (25°C, 20 minutes). Cisplatin (Cat. No.: PHR1624) obtained from Merck™ was dissolved in Milli-Q water (18.2 MΩ cm distilled deionized water obtained from a Milli-Q system, Millipore™, Bedford, MA, USA), forming a cisplatin solution.

### 4. Survival Analysis

All Drosophila survivors were used in survival analysis. Afterward, a phenotypic analysis was conducted using a stereoscope to assess any phenotypic alterations resulting from the treatments. The Kaplan-Meier method was used to calculate the survival rate, followed by the Log-rank (Mantel-Cox) test. Survival rate was performed in R Studio using the “survival package”.

### 5. Locomotion assay

The locomotion assay was performed on the surviving flies after the survival rate had been calculated. The flies were placed in a vertical glass tube (length: 15 cm; diameter: 2.5 cm) marked at 10 cm. After 1 minute, the flies that crossed the 10 cm mark and those that remained at the bottom were counted separately. Three trials were conducted per experiment, with 1-minute intervals between trials. The values obtained represent the mean number of flies that crossed the mark (Top) and those that remained at the bottom (Bottom). Results are presented as the mean ± standard error of the mean (SEM) of values obtained from three independent experiments. For each experiment, a performance index (PI) was calculated, defined as: 0.5[(nTotal flies + Top - Bottom)/nTotal flies]. We performed statistical analysis of the performance indices using Fisher’s test with Bonferroni correction, and we conducted analysis of variance (ANOVA).

### 6. RNA isolation

We used two female flies from each treated group (CIS, NPTIO2, and CIS/NPTIO2) and the control group for RNA isolation. Flies were placed in 1.5 ml RNase-free plastic tubes containing 500 µl of Trizol™ reagent (ThermoFisher™ Scientific, Waltham, Massachusetts, USA) and macerated. Total RNA extraction was made according to the manufacturer’s instructions. At the end of the extraction, the pellet containing RNA was resuspended in 20 mL of water.

To avoid DNA contamination, we used the RNase-Free DNase Set (QIAGEN, Cat. No.: 79254) according to the manufacturer’s instructions. RNA integrity was analyzed using agarose gel visualized in Molecular Imager_®_ Gel Doctm XR+ Imaging system (Bio-Rad Laboratories, Inc.) and quantified with NanoDrop 2000 spectrophotometer (ThermoFisher™ Scientific) and a Qubit 3.0 Fluorometer (ThermoFisher™ Scientific).

### 7. RNA-seq

RNA-Seq library preparation and sequencing were performed at Computational Genomics Unity Darcy Fontoura de Almeida (UGCDFA) of the National Laboratory of Scientific Computation (LNCC) (Petrópolis, RJ, Brazil). The mRNA sequencing libraries were constructed using the Illumina Stranded mRNA Prep Ligation kit (Illumina, San Diego, CA, USA), according to the manufacturer’s protocol. The quality of libraries was verified using the Agilent TapeStation 4200 System (Agilent Technologies, USA) according to the manufacturer’s instructions. Sequencing was performed on an Illumina NextSeq 500 platform (Illumina, San Diego, CA, USA) using the NextSeq 500/550 high-output kit v2.5 (150 cycles) (Illumina, USA), generating 2 × 75-bp reads.

### 8. Analysis of RNA-Seq data

The quality of the raw sequence data was analyzed using the FastQC software (http://www.bioinformatics.babraham.ac.uk/projects/fastqc). Adapter sequences and low-quality regions were removed by trimming the first five nucleotides from the 5’ end and the last 40 nucleotides from the 3’ end of each replicate. Reads shorter than 20 nucleotides after trimming were removed using the Trimmomatic tool (Bolger; Lohse; Usadel, 2014). The remaining readings were aligned to the reference genome of *Drosophila melanogaster* ASM2977511v1, date: April 17_th_, 2023 (BioProject, NCBI), using the BWA aligner (Li; Durbin, 2009). The alignment quality was evaluated using Samtools (Li *et al*., 2009), and the number of reads mapped to each gene was quantified using HT-seq Count tool (Anders; Pyl; Huber, 2015).

### 9. Statistical analysis

DESeq2 (Love; Huber; and Anders, 2014) was used for differential gene expression analysis. Differentially expressed genes (DEGs) were identified using a log2 fold change (log2FC) cutoff > 1 and an adjusted p-value (padj) < 0.05. All treated groups were compared against the control group.

Raw data are available at Gene Expression Omnibus website with GEO Submission number GSE334628.

### 10. Downstream analysis

Functional enrichment analysis of DEGs was performed using the databases: FlyBase (Öztürk-Çolak *et al*., 2024), FlyEnrichr (Kuleshov *et al*., 2019), STRING database (Szklarczyk *et al*., 2023) and Gene Ontology (GO) (Ashburner *et al*., 2000; The Gene Ontology Consortium *et al*., 2026).

## RESULTS AND *DISCUSSION*

### Data analysis

To evaluate the individual and combined *in vivo* effects of cisplatin (50 μg/mL) and TiO_2_ nanoparticles (50 μg/mL), *we measured Drosophila melanogaster* mortality rates as described in the Materials and Methods section. As illustrated in Figure 1, no significant variation was observed among the evaluated groups (p = 0.76).

**Figure 1:**
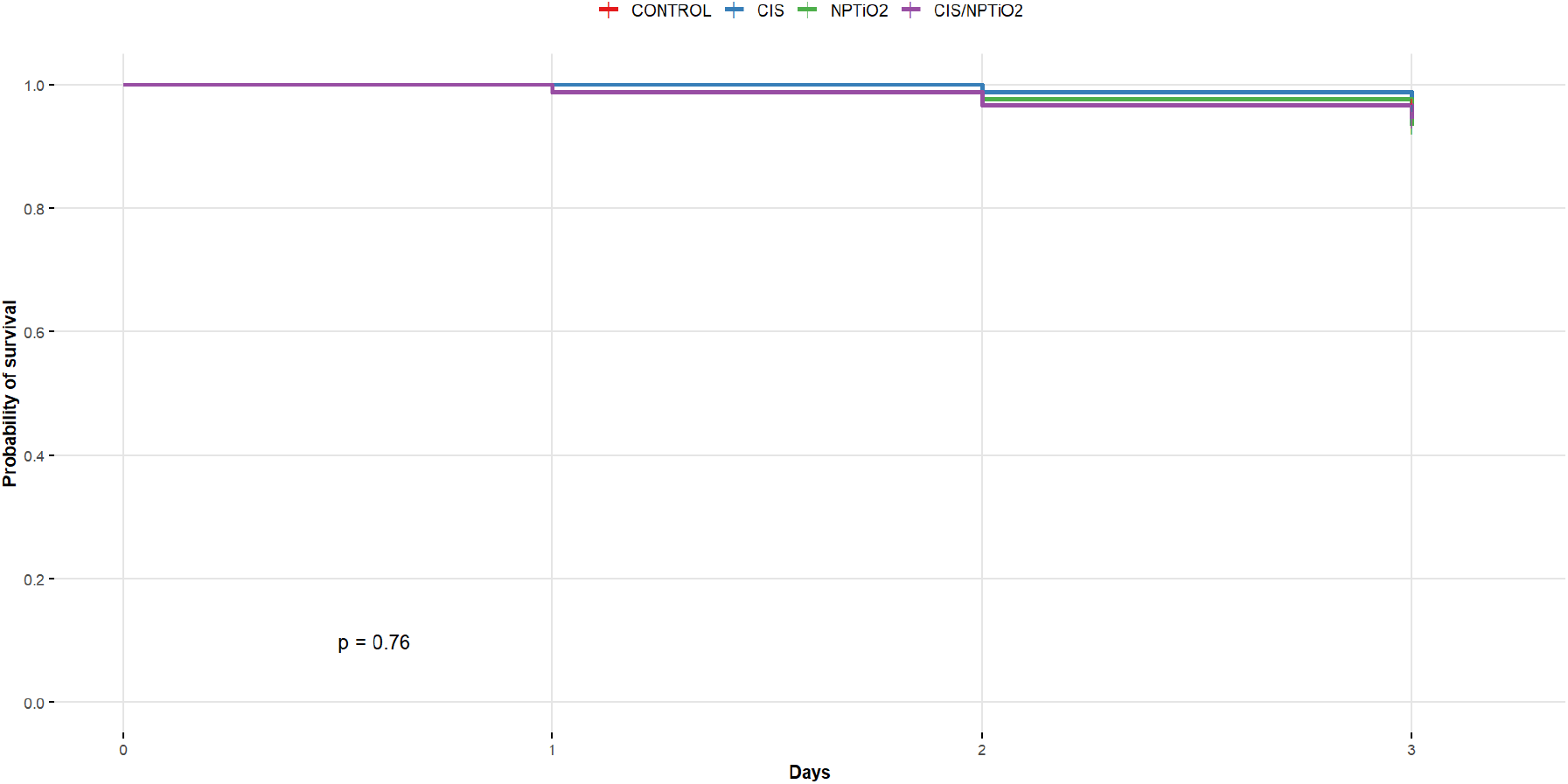
Kaplan–Meier survival curves of *Drosophila melanogaster* exposed to different treatments. Control, cisplatin (50 μg/ml CIS), titanium dioxide nanoparticles (50 μg/ml NPTIO2), and the combination CIS/NPTIO2 (50 μg/ml/50 μg/ml). Survival rates were similar across all experimental groups (p = 0.76). The Log-rank (Mantel–Cox) test showed no statistically significant differences among the groups (Events: 18; p-value: 0.75347), indicating that none of the treatments significantly affected fly survival over the 3-day observation period.

We also performed a locomotion test to assess potential neurological damage in the surviving Drosophila after exposure to cisplatin (50μg/ml) and NPTiO2 (50μg/ml) isolated or in combination (Fauzi; Zubaidah; Susanto, 2020; Mishra *et al*., 2019), the resulting data is presented in Figure 2 and indicates that flyes from CIS/NPTIO2 group exhibiting the lowest proportion of flies reaching the top of vial (62%), which was significantly lower than that of the control (p = 0.041), CIS (p = 0.009), and NPTIO2 (p = 0.001) groups. These results suggest a negative neurological effect of the combined treatment of CIS and NPTIO2.

**Figure 2:**
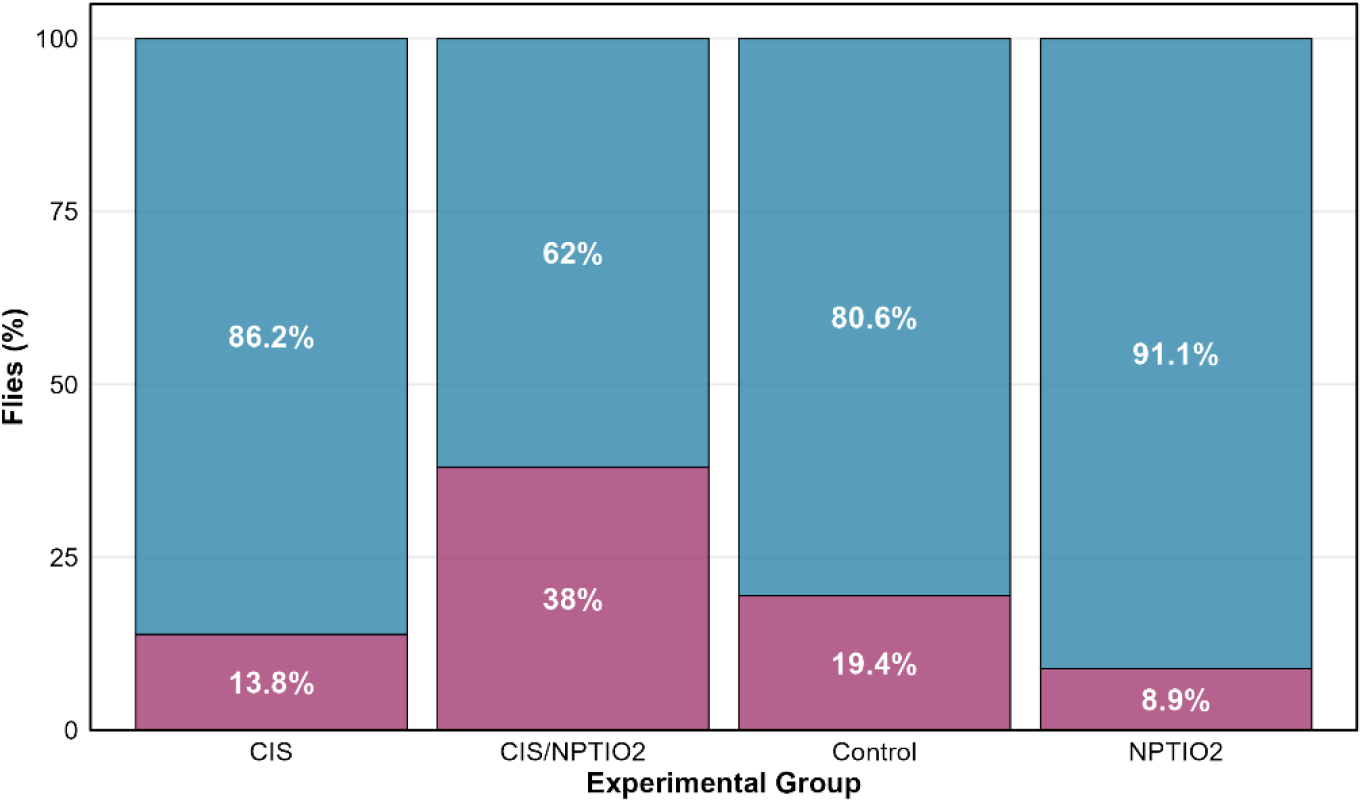
Locomotion assay in *Drosophila melanogaster*: Bars represent the percentage of flies that reached the top (▪) of the vial (mean ± 95% CI) and those that remained at the bottom of the vial (▪). Four experimental groups were tested: Control (n=72), CIS (n=65), NPTIO2 (n=45), and CIS/NPTIO2 (n=50).

To evaluate sample expression variability following DESeq2, we conducted a Principal Component Analysis (PCA) using the BiocManager package. Within this framework, the standard PCA executed by the DESeq2 package applies a regularized logarithm (rlog) transformation for data normalization. PCA is a widely accepted method for variance analysis. As shown in Figure 3, this analysis suggests that CIS/NPTIO22 represents an outlier sample.

**Figure 3.**
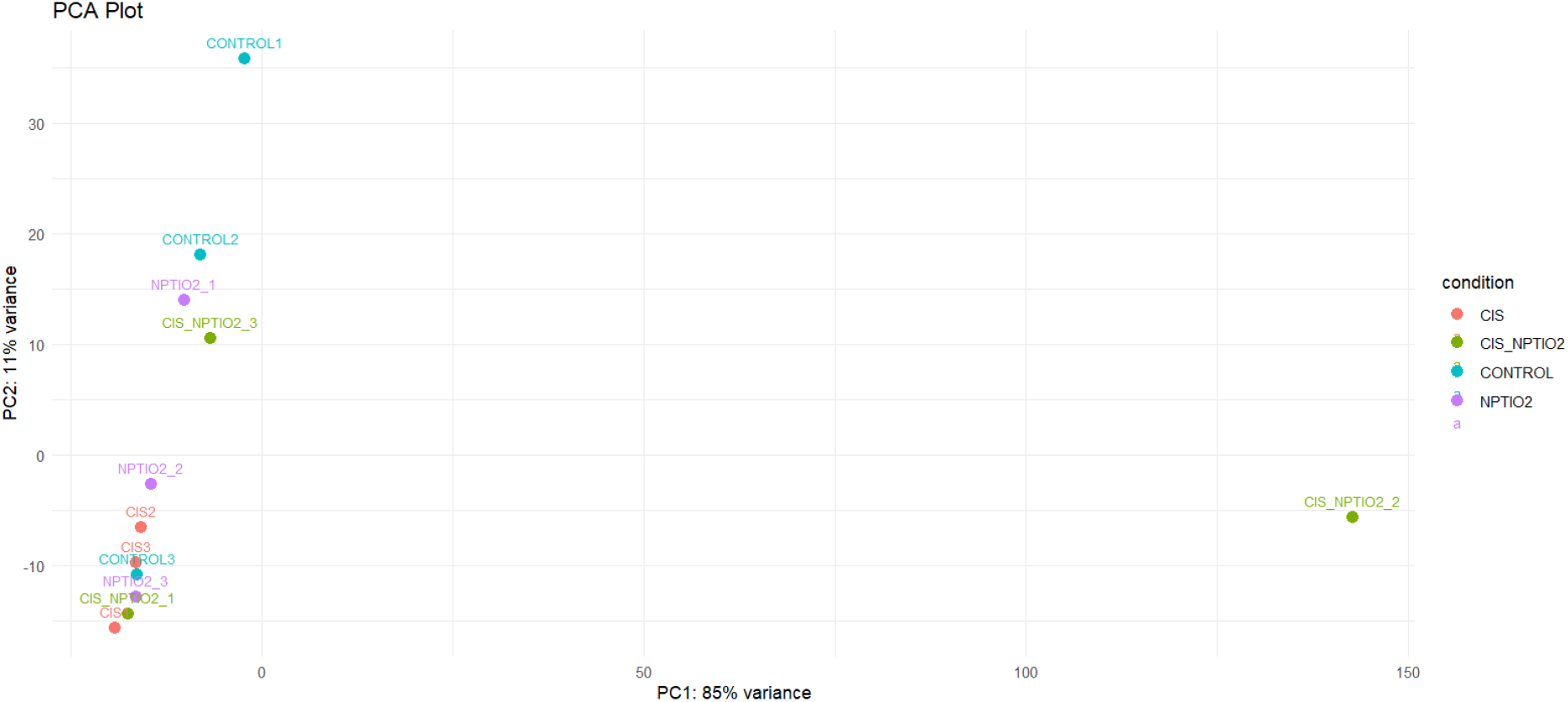
PCA plot of RNA-seq data (rlog-normalized) showing sample separation by treatment. Controls (blue) cluster on the left, while CIS (red), CIS/NPTIO2 (green), and NPTIO2 (purple) form distinct groups along PC1 (85% variance). PC2 explains minimal variation (11%). CIS/NPTIO22 exhibits an intermediate profile between CIS and NPTIO2.

The rlog applies a variance-stabilizing log transformation tailored to count data and prioritizes biological interpretability. Robust Scaler is another normalization method that standardizes data using the median and interquartile range (IQR), making it resistant to outliers and generally applicable (not RNA-seq-specific), with a primary focus on statistical robustness. To optimize PCA results by balancing variance stabilization and outlier resilience, we performed a PCA analysis that combines both methods (rlog followed by Robust scaling).

Here, we analyzed the similarity of expression patterns across the top 10,000 most differentially expressed genes (DEGs) in each treated group after robust scale analysis, comparing within the CONTROL group (lower p-values). Then, we performed a quantitative analysis to verify the Euclidean distances among the replicates, as shown in Table 1, and between CIS/NPTIO22 and all other samples. This data suggest that CIS/NPTIO22 is an outlier.

**Table 1:** Euclidean Distance between CIS/NPTIO22 sample and other samples. Scale units in standard deviation (Robust Scaler).

| CIS1 | CIS2 | CIS3 | CIS/NPTIO2<br>1 | CIS/NPTIO2<br>3 |  |
| --- | --- | --- | --- | --- | --- |
| **4974.11** | 4949.75 | 4944.15 | 4950.60 | 4940.42 |  |
| CONTROL1 | **CONTROL2** | **CONTROL3** | **NPTIO21** | **NPTIO22** | **NPTIO23** |
| **4919.46** | 4926.30 | 4950.49 | 4948.30 | 4943.52 | 4933.49 |

The two-dimensional PCA (Figure 4) showed that CIS/NPTIO22 has a distance greater than 4000 Standard Deviations (SD) from other CIS/NPTIO2 replicates, and similar distances are observed for all samples from other treatment groups. These data are consistent with the outlier condition of CIS/NPTIO22, indicating that this sample should be excluded from further analysis to maintain data integrity.

**Figure 4.**
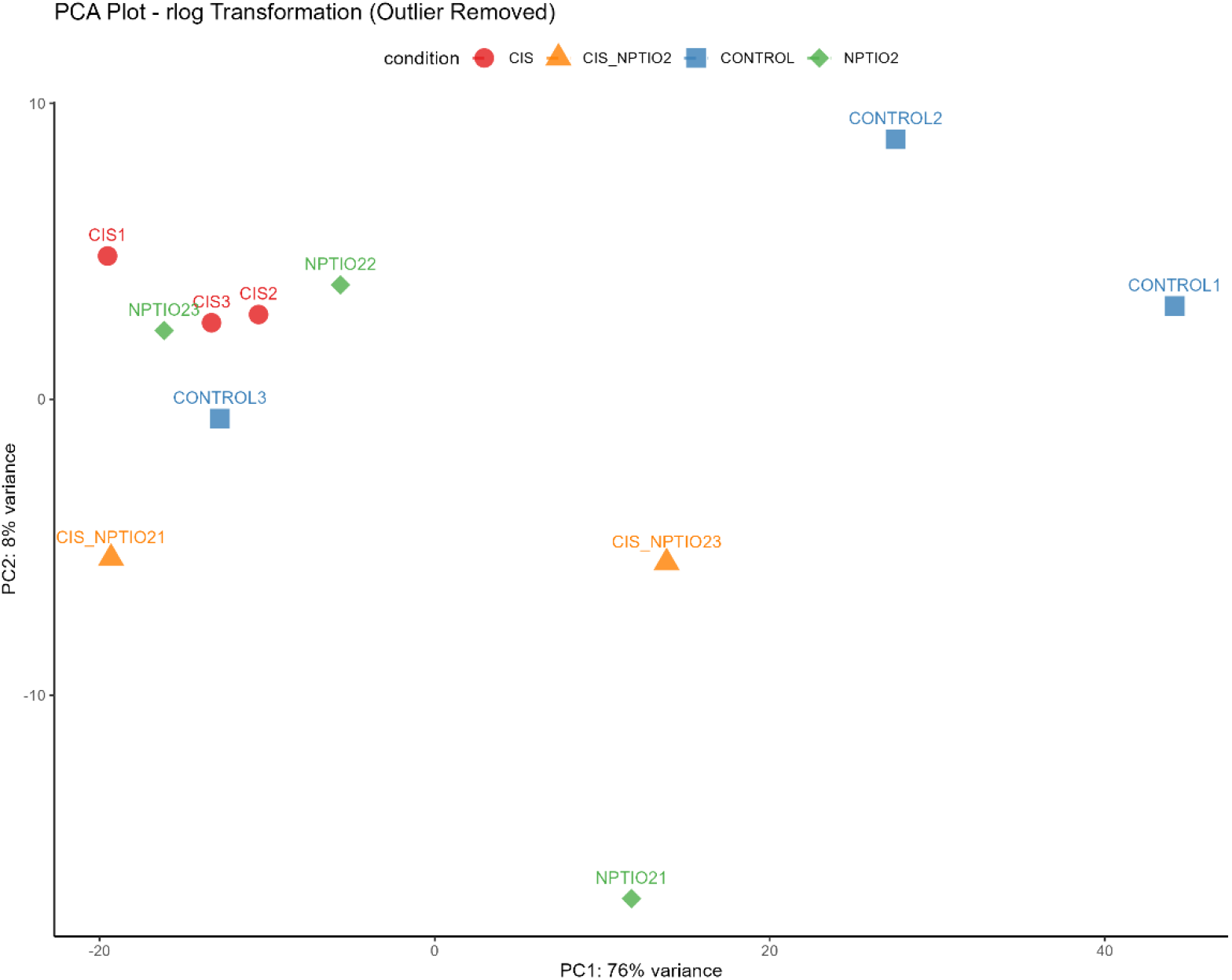
Two-dimensional PCA projection of rlog-transformed count data following quality control and outlier removal. Samples are colored by experimental conditions: CONTROL (blue), CIS (red), CIS/NPTIO2 (orange), and NPTIO2 (green). Variance explained: PC1 (76%), PC2 (8%).

After outlier removal, we repeated PCA and quantitative analysis to assess Euclidean distances among replicates. As shown in Figure 4, PC1 and PC2 explain 70% and 8% of the total variance, respectively. The distances among samples along PC2 indicate only minor biological variation within groups. Nevertheless, along the PC1 axis, CONTROL is the most variable group. CONTROL3 appears more distant (approximately 40%) from CONTROL1 and CONTROL2, though still within the range of expected biological variability. Despite this heterogeneity, the control group. Despite this heterogeneity, the control group remains useful for comparing transcriptional variance across all treatment groups. The CIS group was the most homogeneous, since its samples are closer to each other in both components.

Besides that, PC1 shows heterogeneous responses across samples within treatment groups. NPTIO21 is separated from NPTIO22 and NPTIO23 in both PC1 and PC2, whereas CIS/NPTIO21 is separated from CIS/NPTIO23 only in PC1. These results demonstrate biological variation among samples within those groups, but the variation is not sufficient to discard any of the samples. PC1 explains the largest proportion of variance and primarily separates the treatment groups (CIS, NPTIO2, and CIS/NPTIO2) from the control group, indicating a treatment effect.

To verify the absence of additional outliers, we computed Euclidean distances among the replicates after removing the initial outliers. Neither the control nor the treated group has a distance greater than 750 from other replicates, indicating no further outliers (*Supplemental material 01*).

Taken together, Figure 4 and Table 1 demonstrate that removing the CIS/NPTIO22 outlier yields greater consistency across biological replicates and supports robust inference about treatment effects, enabling more reliable downstream comparisons and interpretation.

### Gene Expression and Downstream Analysis

Building on the above analysis, we next examined global patterns of gene expression and differential regulation elicited by each treatment group.

Volcano plots in Figure 5 compare total transcript counts across treatments, showing the distribution of DEGs (fold change (log2fc) > |1.0| and p-value < 0.05) for each treated group (CIS, NPTIO2, and CIS/NPTIO2) relative to the control.

**Figure 5.**
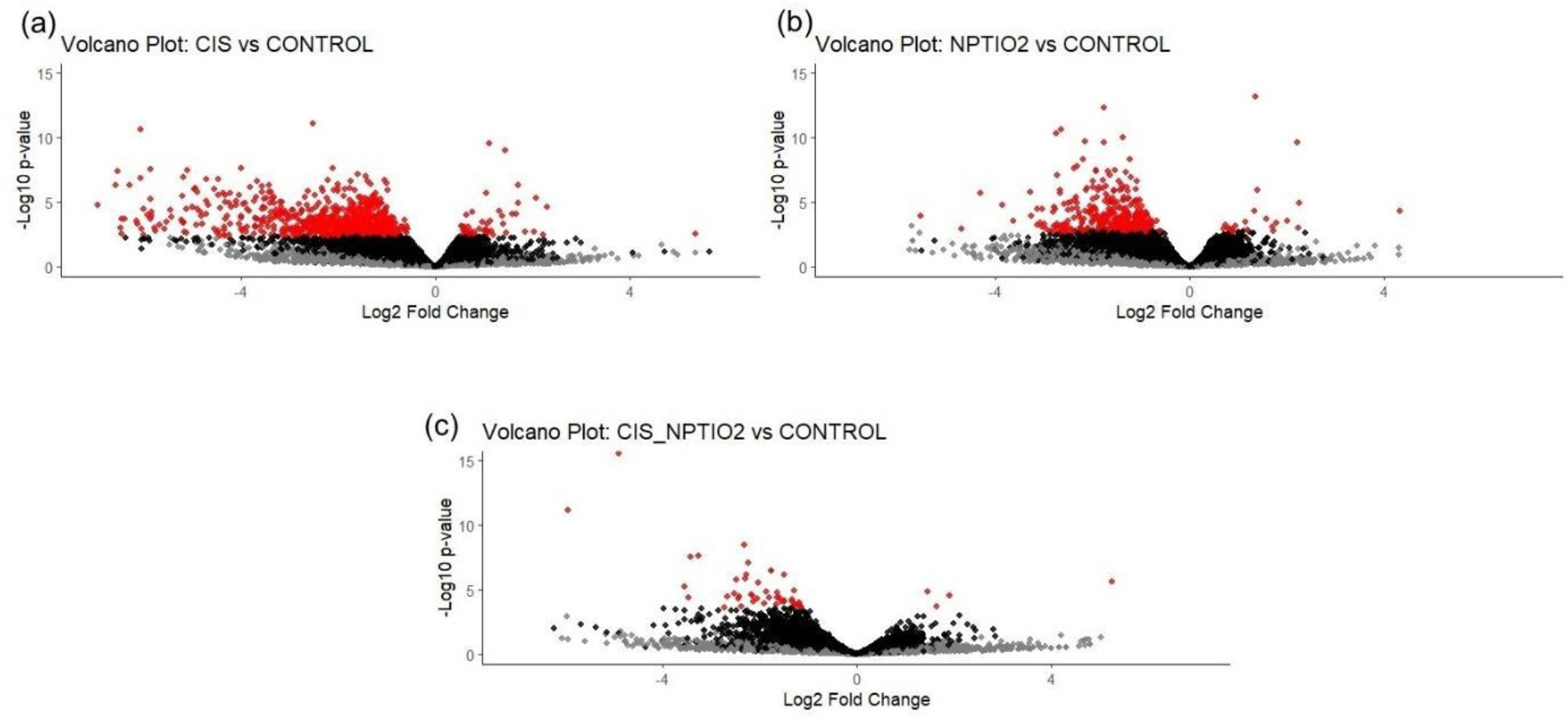
Differential gene expression analysis. Volcano plots display the relationship between statistical significance (-log₁₀ p-value) and the magnitude of change (log_2_ fold change) for all measured genes. (a) CIS treatment versus CONTROL, (b) NPTIO2 treatment versus CONTROL, (c) CIS/NPTIO2 treatment versus CONTROL. DEGS with differentiated expression (fold change (log2fc > |1.0| and p-Value <0.05): Significant (•), Not significant (•).

The total transcript counts by each control or treatment group are indicated in Figure 6. Compared with CONTROL, our data show that both CIS and CIS/NPTIO2 have higher median total transcript counts than CONTROL. However, NPTIO2 treatment alone appears to have a lower total transcript count compared with other treatment groups. As shown in Figure 6, these treatments (particularly those involving CIS) increased transcriptional activity relative to the control baseline.

**Figure 6.**
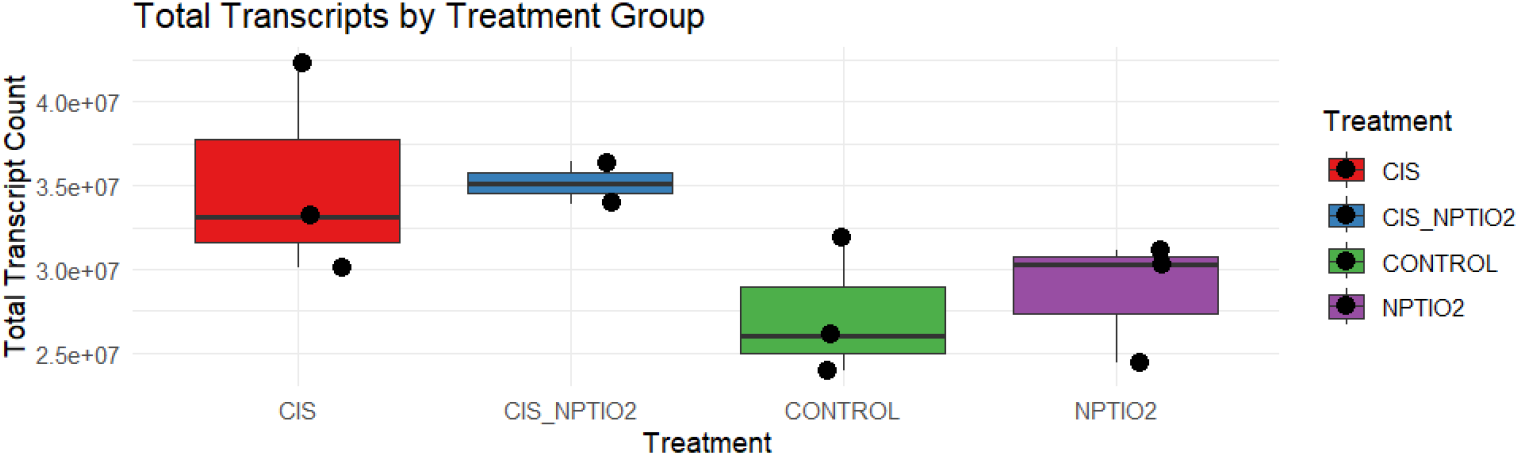
Total Transcript Counts by Treatment Group: Box plots for each of the four treatment groups (CIS, CIS/NPTIO2, CONTROL, and NPTIO2). The line within each box indicates the median, and the box represents the interquartile range (IQR). Black dots represent individual sample data points. Total transcript counts are highest in the CIS and CIS/NPTIO2 treatment groups, while the CONTROL group shows the lowest counts.

Although the CIS/NPTIO2 group presented a lower number of upregulated and downregulated DEGs, the position of its median and interquartile range indicates that the combination of cisplatin (50 μg/ml) and TiO_2_ nanoparticles (50 μg/ml) is effective in keeping the *Drosophila’s* global transcription machinery in a state of hyperactivity.

Figure 7 displays a Venn diagram showing the numbers of uniquely upregulated and downregulated DEGs per group, alongside the shared DEGs found in two or more groups. We detected 28 upregulated and 890 downregulated DEGS in the CIS group, 17 upregulated and 400 downregulated DEGs in the NPTIO2 group, and only 4 upregulated and 61 downregulated DEGs in the CIS/NPTIO2 group. The number of downregulated DEGs in the CIS/NPTIO2 group indicates that both cisplatin and TiO2 have a repressor effect on gene expression in this group.

**Figure 7:**
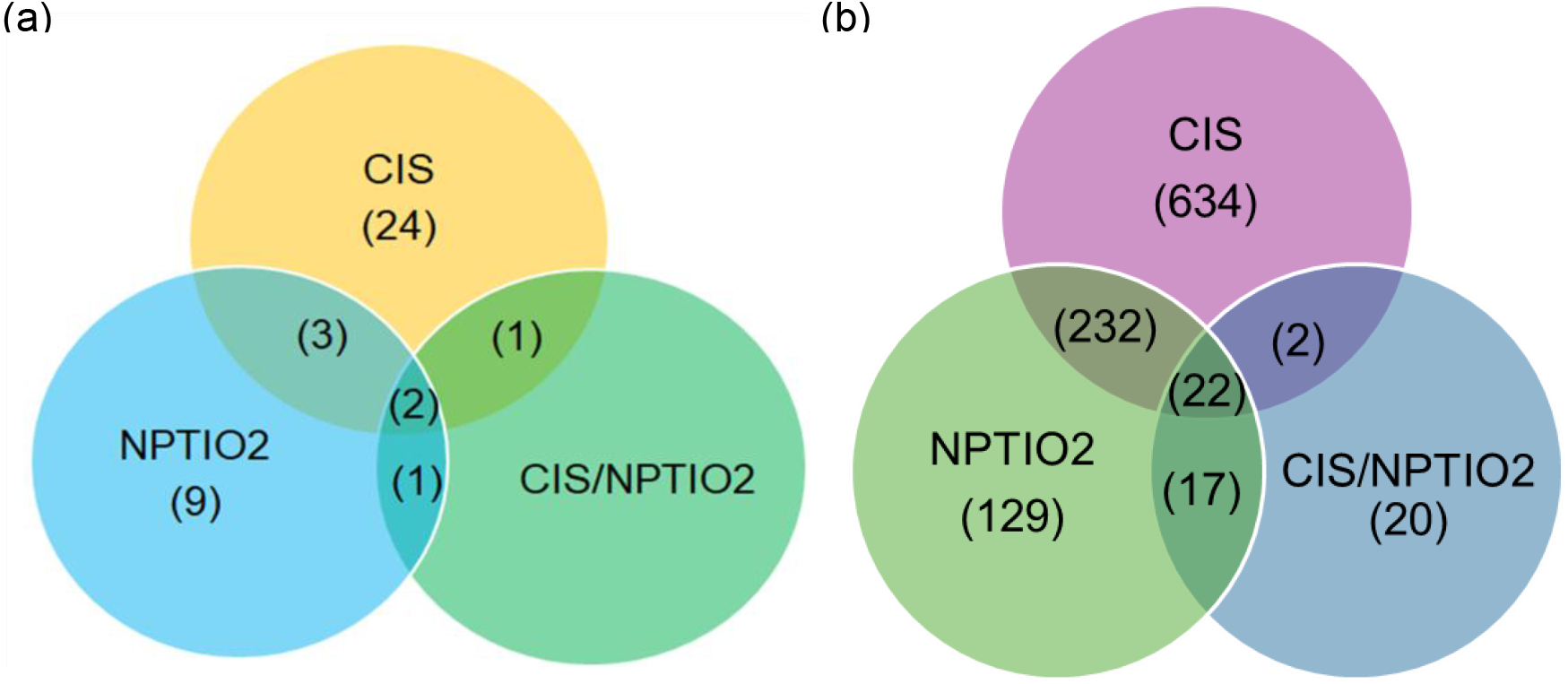
Venn diagram relative to the number of DEGs obtained in (a) upregulated DEGs and (b) downregulated DEGs in the challenged groups.

Two DEGs are upregulated across all groups (CG10013 and aqz). The CIS and NPTIO2 groups share 4 upregulated DEGs (CG11263, CG18088, Tango6, inaF-D) and one long non-coding RNA (lncRNA: CR43836). One upregulated DEG is present in the CIS group and also in the CIS/NPTIO2 group (CG5568), and one upregulated DEG (CG3213) is also upregulated in the NPTIO2 group and the CIS/NPTIO2 group. Conversely, treatment conditions yield distinct profiles of downregulated DEGs across groups. Approximately 19,000 genes were differentially expressed, of which only 1,056 were downregulated DEGs. Among these, 22 DEGs were consistently downregulated across all treated groups. The CIS group had the highest number of downregulated DEGs (890), whereas the CIS/NPTIO2 group had only 61.

The reduced number of up- and downregulated DEGs, particularly in the CIS/NPTIO2 group, suggests a possible synergistic effect of co-administering cisplatin and TiO2 nanoparticles in Drosophila. These results likely reflect reduced cell signaling following co-administration of these substances in Drosophila, as previously suggested by(Adibzadeh *et al*., 2021).

CIS/NPTIO2 groups are primarily elicited by cisplatin exposure. CG5568, annotated as a CoA-ligase/fatty acid ligase, contains AMP-binding enzyme domains (Pfam PF00501 and PF13193), characteristic of acyl-CoA synthetases involved in fatty acid. To assign biological significance to the above-mentioned upregulated genes, we performed an in silico functional analysis using STRING database annotations, focusing on protein domains, Gene Ontology (GO) terms, subcellular localization, and associated mutant phenotypes.

We identified two genes upregulated across all three treatment groups: CG10013 and aqz. CG10013 is largely uncharacterized and encodes a ubiquitin-conjugating enzyme E2-binding protein (Pfam PF09814) that accepts ubiquitin from specific E2 ubiquitin-conjugating enzymes and transfers it to substrates, generally promoting their degradation by the proteasome. CG10013 has potential involvement in pigmentation variation, alongside other genes known to be involved in melanin production, such as ebony, tan, yellow, and black (Bastide *et al*., 2016). The gene aqz is predicted to be active in the nucleus and codes for the Aaquetzalli protein isoform A. aqz is orthologous to human PNRC1 (proline-rich nuclear receptor coactivator 1) and PNRC2 (proline-rich nuclear receptor coactivator 2) (Pfam PF15365, InterPro IPR026780) and is related to neural tissue formation and nervous cells polarity there is no known function of aqz in adult flies (Mendoza-Ortíz; Murillo-Maldonado; Riesgo-Escovar, 2018). However, our data indicate that together, CG10013 and aqz play a role in responses to oxidative stress and other conditions affecting the cell, as evidenced by their upregulation across all treated groups.

The upregulation of CG5568 was observed in flies from both the CIS and β-oxidation and complex lipid biosynthesis. Its association with lethality and P-stage mortality suggests an essential role in maintaining energy homeostasis during late embryonic or larval development; also, this gene is highly expressed in female germline cells and implicated in the fatty-acyl-CoA biosynthetic process (Li *et al*., 2022a, 2022b). Our findings about the shared lethal phenotypes of aqz and CG5568 in embryonic and larval stages point to a potential convergence between nuclear receptor-mediated transcriptional control (via aqz) and metabolic regulation (via CG5568), which could explain the known side effects of cisplatin in adult flies.

CG3213 is upregulated in flies from both NPTIO2 and CIS/NPTIO2 groups; this gene is involved in ribosomal and protein translation (Pospisilik *et al*., 2010); it is still predicted to regulate organelle organization and cilium assembly, with strong localization signals for the centrosome and microtubule cytoskeleton. Its mutant phenotypes, including abnormal pain and stress responses, are consistent with a role in neuronal or sensory cell architecture. CG3213 mutant phenotypes include abnormal pain and stress responses, indicating a role in neuronal or sensory cell architecture, where microtubule-based transport and centrosomal positioning are paramount.

Based on their predicted molecular functions and phenotypic data, allowed us to infer that CG10013 and CG3213 may act by modulating the stability of proteins involved in microtubule organization, processes that depend on both vesicular trafficking and cytoskeletal integrity, which may explain the absorption of metal particles, such as cisplatin and TiO2 nanoparticles, by the adult Drosophila.

The above observation, combined with the biological processes enrichment analysis of the upregulated DEGs of CIS/NPTIO2 described in Table 2, specifically those related to fatty-acid and lipid metabolism, strongly suggests a synergistic effect of the cisplatin and TiO2 nanoparticles jointly administered in D.melanogaster due to a possible increase in the incorporation of cisplatin due to the endocytosis mediated by the NPTiO2 joint administration.

**Table 2:** Gene Ontology (GO) enrichment analysis of CIS/NPTIO2-upregulated genes.

| GO Category | Subcategory | Nº Genes |
| --- | --- | --- |
| Cellular Component | Cytoskeleton | 1 |
|  | Nucleus | 1 |
| Biological Process | fatty-acyl-CoA biosynthetic process | 1 |
|  | long-chain fatty acid metabolic process | 1 |
|  | biological_process | 1 |
| Molecular Function | Enzyme | 1 |

Drosophila from both CIS and NPTIO2 groups have 3 upregulated DEGs in common (CG11263, CG18088 and InaF-D) and an upregulated long non-coding RNA (lncRNA:CR43836). Recent data suggest that long non-coding RNAs (lncRNAs) may have important regulatory functions in the genome (Li *et al*., 2019). Specifically regarding lncRNA:CR43836, data show that lncRNA:CR43836 is expressed in many embryonic tissues, larval visceral muscle cells, and adult females (Matthews *et al*., 2015).

CG11263 is involved in RNA-related regulation, including piRNA metabolic process, gene silencing by RNA, germ plasm/P granule localization, and is expressed in Drosophila embryos, prepupa and pupa stages, and in adult females (Adams *et al*., 2000). Upregulation of CG11263 could explain the presence of lncRNAs in treated flies in these groups, given its role in transcriptional regulation.

CG18088 has reproduction-associated features and is associated with membrane/transmembrane features; it is associated with Drosophila’s immune response to Wolbachia sp. and affects transduction-related genes. InaF-D also has a membrane feature, related to membrane potential regulation and channel regulator activity; it is membrane/transmembrane-associated, binds to the transient receptor potential (TRP) calcium channel (PFAM term: PF15018), and affects metal absorption (Chen; Montell, 2020). The last DEGs are also related to immune and metal response and are expected to be upregulated in flies treated with those metal compounds (López-Varea *et al*., 2021). We propose that the upregulation of these DEGs is due to the absorption of cisplatin and TiO2 NPs.

In this study, all flies were kept for 3 days and fed food supplemented with wet yeast paste containing cisplatin, NPTiO2, or both substances, without prior starvation. This can positively affect cell proliferation in flies, regardless of group (control or treatment) (Ables *et al*., 2016; Ables; Laws; Drummond-Barbosa, 2012). We detected no upregulated DEGs related to oogenesis. In this study, we chose the Oregon strain, which displays standard metabolism that facilitates observation of general metabolic responses to substance administration and toxicity, unlike tumor-bearing strains, which exhibit tumor-characteristic metabolic pathways that could mask the general toxicity response to each treatment.

Previous studies have demonstrated that mated females can store sperm transcripts in their spermatheca, and RNAs related to sperm from many genes, including those of accessory gland and seminal fluid proteins, can be detected (Cridland; Begun, 2023). Our treatment conditions, feeding time, food supplementation provided to flies, use of mated females, and the pipeline and sequencing technologies used could explain the difference between our results and those of the previous study in flies treated with the same cisplatin concentration (Mombach *et al*., 2022). It could also explain the identification of DEGs primarily related to Drosophila males but also expressed in females in our data.

CIS and NPTIO2 groups presented a response whose enrichment has the GO terms (Figures 7 and 8) related to response to stimulus, gene expression, protein metabolism and development (BP); and enzyme, nucleic acid binding and small molecule binding (MF); DEGs in these groups are located in different cellular components but enrichment GO terms related to the membrane, endomembrane system, and macromolecular complex has found in both. Our data are partially consistent with a previous study indicating that, regardless of the treatment to which Drosophila are subjected, upregulated genes have GO terms related to stimulus response and include genes encoding lysozymes, cytochrome, and mitochondrial components (Brown *et al*., 2014).

Our data show upregulated DEGs related to different response-to-stimulus GO terms. According to GO Summary Ribbons (FlyBase (FB2025_03, released July 24, 2025), the CIS group has six upregulated DEGs with Response to stimulus GO Term: FANCI, St4, Ir7c, PPO1, out, and CG10738. A more detailed analysis of the FlyEnrichr database indicates that this set of DEGs is associated with the following GO biological processes, including defense response to bacteria, oogenesis, regulation of cell proliferation, and wound healing, among others (*Supplementary Table 2*). The molecular functions of this set of DEGs are related to sulfotransferase activity, DNA polymerase binding, and hydrolase (*Supplementary Table 2*).

**Table 2**: DEGs from CIS/NPTIO2 group classified using GO Summary Ribbon (Fly Base). Only the most specific terms are shown. Full data available in *Supplementary Table 1*.

CIS group flies also present three upregulated lncRNA (lncRNA:CR43836, lncRNA:CR44707, lncRNA: CR45052). lncRNA:CR43836, as mentioned before, and all knowledge regarding these lncRNAs is related to regulation in transcription. Our data are consistent with the state of the art, which indicates that cisplatin treatment induces lncRNAs-mediated responses (Brown *et al*., 2014; Zhang *et al*., 2020).

The CIS group also has DEGS related to development: CG9503, dec, Cad88C, Cad74A, and out. The FlyEnrichr database shows that this set of DEGS is associated with the following GO biological processes: cell-cell adhesion, female germline sex determination, regulation of DNA endoreduplication, and ecdysteroid metabolic process, among others (*Supplementary Table 1*). These molecular functions are related to hydrolase activity and ionic binding (*Supplementary Table 2*). The combined super expression of all these genes suggests that the organism is undergoing an intense phase of development, differentiation, and tissue reorganization. They appear to be working together to coordinate the complex morphological changes that occur in the Drosophila body in response to cisplatin.

In the NPTIO2 group, we found out that there was upregulation in CG1503, l(2)03659, and CG44838. This upregulation suggests that Drosophila treated with 50 μg/ml TiO_2_ needs to activate transport machinery to metabolize TiO_2_. In case, CG44838 is a transcription factor known for signal transduction and nervous system development (Olson *et al*., 2019) and activate CG1503 which is a protein regulator having hydrolase activity and are related to decreased carbohydrate metabolism (Mok *et al*., 2025), finally l(2)03659 which is related to response to xenobiotics or metabolites mitochondrial stress-relief mechanism associated with multidrug resistance (Yeh *et al*., 2017) due to activation in transmembrane transport mechanism.

CG11263, tut, and CG18088 DEGs were upregulated by NPTiO2. These DEGs are associated with the negative regulation of gene expression and the piRNA metabolic process. CG11263 is co-expressed and functions as part of the transcription machinery that generates piRNA precursors from heterochromatin while maintaining suppression of transposon-encoded promoters and enhancers. This is also related to pupal and adult female viability (Matthews *et al*., 2015). CG10013 and ord are co-expressed and promote normal formation and maintenance of the synaptonemal complex and centromere clustering (Das *et al*., 2016). Ord is also essential for proper maintenance of sister-chromatid cohesion in both male and female meiosis. Yet, mutations in this gene cause premature separation of sister chromatids in meiosis I, random segregation in both meiotic divisions, and reduced recombination in female meiosis (Miyazaki; Orr-Weaver, 1990).

Relative to NPTiO2, a study using *D.melanogaster* chronically exposed to suspensions containing 5, 10, 15, and 20 µg/kg NPTiO2 from the embryonic period of the P0 to F3 in the adult stage demonstrates a decrease in the total crawling distance of larvae and the total movement distance of adult males in F3. Results obtained in this study suggest disruption in the Impaired Neuromuscular Junction (NMJ) due to an increased expression level of genes related to response to Lithium-ion Response (List) and neuron cellular homeostasis (Gba1a and hll), and a decrease in gene expression levels of thermosensory behavior gene (Cyp6a17) and axon target recognition gene (frac) (Zhang *et al*., 2023). Our data, are based on prolonged but not chronic exposure and corroborate these observations regarding the reduction in the expression of frac and Cyp6a17.

Based on our observations aqz and CG10013, which are DEGs upregulated by all treated groups (CIS, NPTIO2 and CIS/NPTIO2), should have a significant role in the fly response to cisplatin and TiO2 nanoparticle. The upregulation of these DEGs across all treated groups indicates that ribosomal production and metabolism are affected by these treatments.

Of particular interest is the observation that the 61 downregulated DEGs in CIS/NPTIO2 group, mostly are related to molecular functions (MF) of hydrolase, serine and serine-peptidase activity; the repression of these MFs may lead to a drastic defects in embryonic development, collapse of the innate immune system, reproductive impairment and fertility, loss of resistance to insecticides and xenobiotics, dysregulation of lipid metabolism and phagocytic degradation (Kumar *et al*., 2021; Zhang *et al*., 2020). This may also be related to high embryonic lethality, pupation failure, and severe immunodeficiency in flies from this group (Marra *et al*., 2021; Mulinari; Häcker; Castillejo-López, 2006).

In conclusion, our data suggest a toxic synergistic effect caused by the combined use of cisplatin and NPTiO2, which leads to neurological impairment and metabolic alterations in Drosophila within the CIS/NPTIO2 group. In particular, flies in this group show no statistically significant difference in survival but the worst performance in the climbing test and alterations in cell-division metabolism. Therefore, the combined administration of cisplatin and TiO_2_ nanoparticles may not be considered an alternative cancer treatment when cisplatin alone is ineffective.

## Acknowledgments

We thank Professor Dr. Helena Maria Marcolla Araújo for kindly providing the fly strains. We also acknowledge FAPERJ and CAPES for partial funding support.

